# Electrochemically controlled switching of dyes for enhanced super-resolution optical fluctuation imaging (SOFI)

**DOI:** 10.1101/2024.06.02.597055

**Authors:** Ying Yang, Yuanqing Ma, Richard D. Tilley, J. Justin Gooding

**Author notes:** Correspondence (Y.M.); (J.G.).

## Abstract

In super-resolution optical fluctuation imaging (SOFI), the locations of molecules spaced closer than the diffraction limit of light can be identified through spatial and temporal correlation analysis of the fluorescence intensity fluctuation. Using organic dyes as fluorophore probes, the fast but stochastic switching of the individual dyes is favourable for improving SOFI imaging resolution and speed, especially in the case of high-order cumulant analysis. While in practice, fluorophore switching can be non-uniform, with some fluorophores remaining in ON or OFF state for extended periods. Furthermore, in some cases the overall rate of switching of the fluorophores can be too slow, presenting practical limitations for high-resolution and fast SOFI imaging. In this work, we demonstrate how to overcome these challenges using electrochemical controlled fluorophore switching. The oscillating electrochemical potential setting with high frequency increases the switching rate and reduces the switching heterogeneity of fluorophores. The dye Alexa Fluor 647, applied here as an example, exhibits over 3-fold decrease in average ON time and over 2-fold decrease in switching variance, resulting in significantly improved SOFI image resolution with fewer frames. We demonstrated that this new electrochemically controlled SOFI imaging modality can achieve a SOFI image with ∼130 nm resolution in 2 seconds of acquisition time, and 80 nm resolution in 6 seconds. This advancement enables fast, large area tile-scan super-resolution imaging, which opens the full potential of SOFI imaging.

## Main

Fluorescent light microscopy has been revolutionized by breaking the diffraction barrier of light using a few different strategies that have led to a range of super-resolution techniques. Among them, a class of methods adapt the illumination pattern to improve the resolution of the overall structure, such as stimulated emission depletion (STED) microscopy[1, 2], structured-illumination microscopy (SIM) [3], and reversible saturable optically linear fluorescence transitions (RESOLFTs)[4]. Another class of methods is based on localization of individual fluorescent molecules in time and space to build up a super-resolution image, including stochastic optical reconstruction microscopy (STORM)[5, 6], photo-activated localization microscopy (PALM) [7, 8] and point accumulation for imaging in nanoscale topography (PAINT)[9]. The super-resolution optical fluctuation imaging (SOFI) represents a third class of super-resolution approach that increases image resolution through higher order cumulant analysis of temporal switching of the dyes[10-12]. It takes advantage of the molecule blinking are independent events which gives unique statistical signatures for the molecules. This is because the photons emitted from the same molecule are correlated in time due to photon bunching effect. Due to quantum mechanical transition between different electronic state, the photons emitted from the same molecules displays bright and dark intervals in succeed to each other in the time scale of microseconds, so that the probability of detecting another photon after a given time delay of a first photon been detected is greater than the case of random distribution in this time scale. This ultimately produces the positive autocorrelation[13, 14]. Due to the diffraction of light, the photon emitted from single molecules are spread across multiple pixels at the camera space. As the images of the molecules from the sample project onto the camera, there are pixels that overlay perfectly to the center of the molecules, and pixels located at the edge between the images of multiple molecules, which can appear even brighter than the former pixel. However, the former pixel that receives photons emission from temporal blinking of single emitters gives rise to a more correlated signal than the later pixels receive non-correlating photons from neighboring emitters. By calculating the temporal correlations of each pixel trace, SOFI alters the image contrast by emphasizing the correlated contributions and diminishing the non-correlating signals, which reduce the blurring effect by light diffraction. Spatially, the fluorescence of single molecules forms a gaussian like spot spread over multiple pixels at the image plane. As the correlation amplitude changes quadratically with the brightness of the pixels, the pixels in the center of the emission is weighted more than the pixels near the edge so that the point spread function from single emitters is sharpened in the SOFI image[11], which ultimately increases image resolution and takes the imaging towards super resolution.

Compared with localization based super resolution imaging method like STORM, the image acquisition conditions for SOFI are greatly relaxed[15]. Generally, SOFI can be performed with high emitter density and low signal to noise ratio[10, 15, 16]. For instance, SOFI can operate at signal to noise ratio in the range of 10-20 dB, where the molecule localization in STORM has failed[15]. Additionally, as overlapping emitters can be tolerated in SOFI, it has been demonstrated that a SOFI can generate images with a 2.5 times improvement in resolution over a conventional diffraction limited widefield image from only 20 image frames with highly switching dyes. This is two orders of magnitude less than the number of images required for a typical STORM acquisition[17]. These advantages make SOFI an easily accessible tool for fast, high-throughput imaging[15, 18, 19].

A SOFI image is essentially reconstructed from the statistical parameters that describe the spatial and temporal fluctuations of the fluorescent events among the pixels. Thus, a SOFI image can be generated if the fluorophores switch ON and OFF stochastically and independently of each other. In principle the resolution of SOFI can be improved infinitely by taking higher order cumulants [10]. The cumulant function is closely related to correlation function, where the higher order cumulant can be directly converted from the same order correlation function by removing the cross-correlation terms, so that only the highest correlating signal are considered. This provides the highest resolution of SOFI image. However, there are practical factors that restrict the resolution of a SOFI image generated using higher order cumulants. These are the heterogeneity in the fluorescence intensities and switching dynamics, which cause the appearance of dark pixels and discontinuities in the image structures[10, 20, 21]. In addition, the overall switching rate of the dyes is usually not fast enough to be comparable to the frame rate, limiting the SOFI imaging speed[21]. Therefore, effective control of the fluorophore switching speed and uniformly is required for SOFI imaging.

Recently, we invented an electrochemical strategy to efficiently switch the organic dyes for STORM imaging, which we refer to as EC-STORM[22]. By tuning the applied electrochemical potentials, the exquisite control over the switching kinetics of the fluorophore was possible which also allowed the emitter density can be effectively altered over a much broader range and with greater control than is achievable with photochemical switching of fluorophores. The efficient ON- and OFF-switching of the dyes using electrochemical potential gives us a hint that we could modulate the dye switching using electrochemical approach to meet the above-mentioned requirements for high-resolution and fast SOFI imaging. Hence, in the present work, we designed an approach to switch the dyes using an oscillating electrochemical potential. The results showed a greater fraction of the dyes in the sample were forced into ON- and OFF-switching cycles. The switching rate of the dyes could be increased by 3-fold, and the variation of dye switching was reduced 2-fold across the sample. Together, these improvements in dye switching resulted in enhancement of SOFI image resolution. We demonstrated that the electrochemical SOFI image modality can be combined with fast sCMOS cameras to achieve 80 nm resolution in ∼6 seconds, and performed large area (0.66 × 0.33 mm) SOFI imaging in 4.5 minutes.

## Results and discussion

### Electrochemical potential modulates the switching of Alexa 647

To achieve stochastic switching of Alexa 647 as required for SOFI imaging, a low oxygen buffer in presence of thiol is utilized, which is also the typical buffer composition for STORM imaging[23]. During the conventional photochemical approach for SOFI imaging, an intense excitation laser is used to switch the dyes to off-states, and the dyes then enter a stochastically blinking mode for SOFI imaging. To facilitate the recovery process of off-state dyes back to on-states, a second short-wavelength laser is normally applied. Here, we replace the photochemical method with electrochemical approach to switch the dyes on and off, as the electrochemical strategy showed higher efficiency and faster switching kinetics demonstrated from the EC-STORM imaging[24]. With the purpose of applying electrochemical potential and microscopy imaging simultaneously, indium tin oxide coated glass (ITO) was chosen as the surface due to its dual property of high conductivity and good transparency[22, 25, 26]. Single Alexa 647 labelled DNA origami samples were used for imaging. To increase the signal to noise ratio of the imaging data, we acquired the image in total internal reflection (TIRF) mode. We first demonstrated that the electrochemical potential could effectively switch the dyes on and off, and thereby tune the density of ON-state Alexa 647 molecules. As Figure 1a shows, the emitter density under ‘no potential’ condition represents the normal emitter density using a conventional photochemical switching method. The results showed that a negative potential of -0.8 V can turn off nearly all the Alexa 647 molecules electrochemically, resulting in a much lower density than no potential was applied. When a positive potential of 0.4 V is applied, many more molecules recover back to fluorescent state, and the emitter density is much higher than under ‘no potential’ condition.

**Figure 1.**
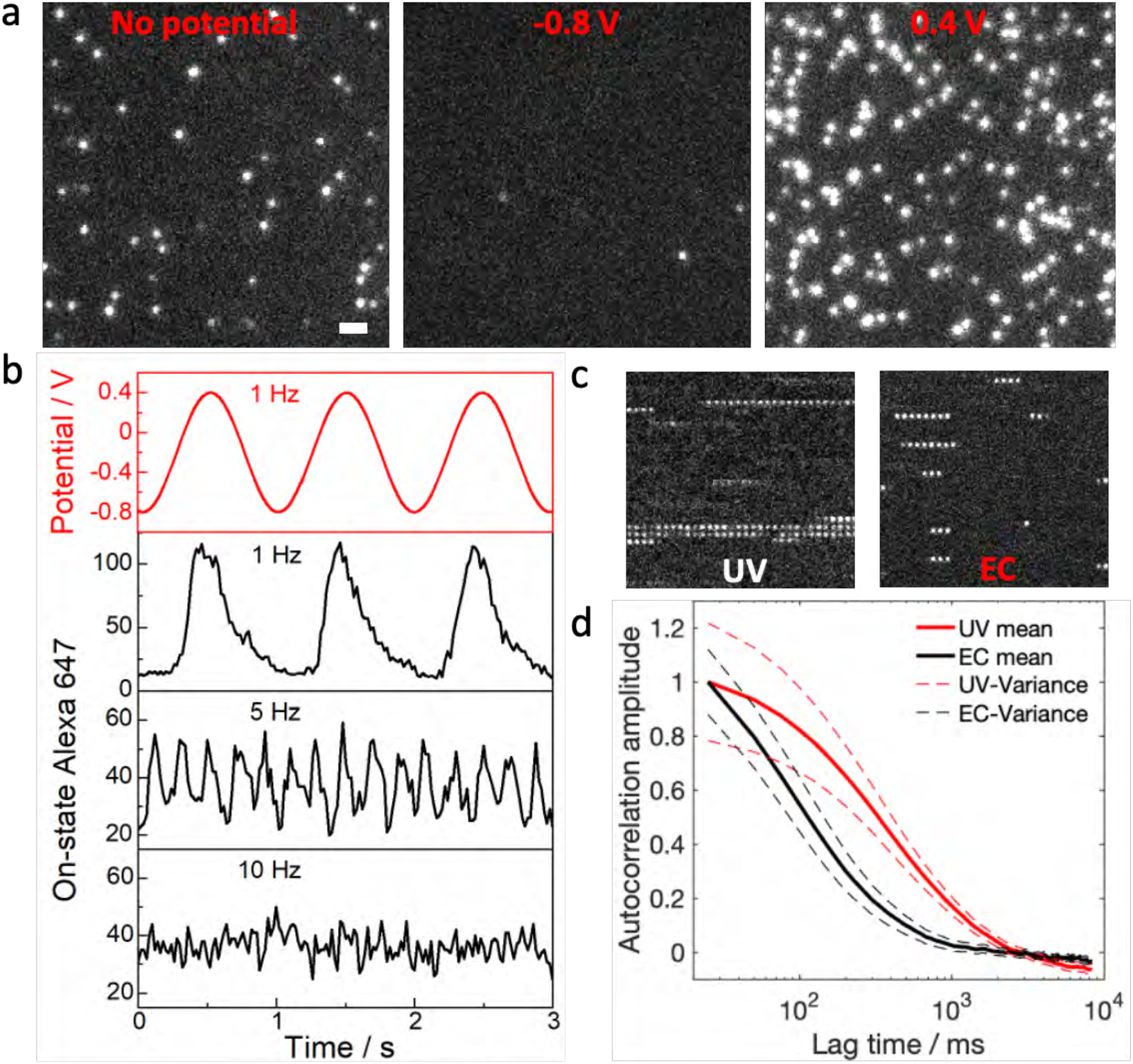
Electrochemical switching of Alexa 647. **a**, TIRF images of ON-state Alexa 647 with no potential applied, by applying electrochemical potential of -0.8 V and 0.4 V, respectively. Scale bar = 2 µm. **b**, The Alexa 647 molecules are turned ON and OFF while applying a sinusoidally oscillating potential between -0.8 V and 0.4 V, as a result, the number of ON-state Alexa 647 in each frame changes with the sinusoidally oscillating potential under various frequencies as indicated. The corresponding Supplementary Movie 1 shows the dynamic switching behavior of Alexa 647 molecules under oscillating potential of 1 Hz. **c**, Example montage of a 1×1 µm^2^ region from time-lapse imaging of single Alexa 647 molecules under UV laser activation condition (UV, 5 W cm^-2^ UV laser is on) or electrochemical potential oscillating condition (EC, the potentials were oscillated between -0.8 V and 0.4 V with a frequency of 10 Hz). The montage was arranged in a raster scan from left to right and top to bottom. **d**, Temporal autocorrelation of Alexa 647 intensity under UV illumination (5 W cm^-2^, blue) or oscillating electrochemical potentials (the potentials were oscillated between -0.8 V and 0.4 V with a frequency of 10 Hz, red). The imaging sample is a DNA origami labelled with a single Alexa 647 labelled, the imaging and electrochemical solution is an oxygen scavenger buffer with 50 mM cysteamine. A 0.5 kW cm^-2^ 642 nm laser is on for imaging.

The capability of electrochemical switching of dyes goes beyond controlling the emitter density. We here intend to enhance the switching kinetics of Alexa 647 by cycling the potential between negative and positive values. To test the feasibility, we firstly oscillated the potential sinusoidally between -0.8 V and 0.4 V at a frequency of 1 Hz or 5 Hz. The results in Figure 1b and Supplementary Movie 1 show the ON-state density of Alexa 647 molecules oscillated with the same frequency as the potential scanning. However, synchronous activation and deactivation of multiple dyes would render SOFI ineffective[10], especially for the dyes that are in adjacent pixels. To minimize the synchronous on and off switching, we then increased the electrochemical oscillation frequency to 10 Hz. At this frequency, the intrinsic stochastic switching of the dyes became dominant, and the overall switching kinetics of dyes were much accelerated. In the case of UV laser activation mode without applying potentials, the ON durations of the organic dyes can sometimes be quite long. As the example showed in Figure 1c, the single Alexa 647 exhibits extreme long ON time, and this lack of fluctuations will eventually give dim signals in SOFI images. With electrochemical switching, single Alexa 647 molecules under the oscillated potential at 10 Hz exhibit much shorter duration for each ON event and undergo more switching cycles in a given time, see Figure 1c. To estimate the switching kinetics of dyes from thousands of Alexa 647 molecules, we conducted autocorrelation analysis of pixel intensity fluctuations in the image series (Figure 1d and Supplementary movie 2). The correlation length provided valuable information about the average ON-time of the fluorophores. Under oscillating electrochemistry modulation at 10 Hz, we observed an over 3-fold decrease in the average ON-time (∼160 ms) of Alexa 647 compared to its ON-time (∼550 ms) during spontaneous switching under UV illumination. This is attributed to the electrochemical switching disrupts the intrinsic switching process of the dyes. Furthermore, we discovered that the spatial heterogeneity of Alexa 647 switching was significantly reduced with electrochemical manipulation. A comparison of the variance of the pixel autocorrelation across the sample revealed a 2-fold reduction with electrochemical control as opposed to the UV laser control at a lag time of 100 ms. This reduction in variance indicates a more consistent and uniform switching behavior of Alexa 647, which would further benefit SOFI imaging by minimizing variation of pixel brightness across the sample. These results highlight that electrochemical control significantly improves two essential imaging conditions for SOFI imaging. Firstly, it accelerates the dye switching kinetics, leading to increased correlation amplitude and pixel brightness in SOFI image. This advance would enable faster SOFI imaging with less frame acquisition [17]. Secondly, electrochemical control ensures consistent dye switching kinetics throughout the sample, and hence reduces spatial and temporal variations to generate more uniform SOFI pixel brightness across the sample[10].

### Oscillated electrochemical potential benefits SOFI imaging

We next conducted experiments to assess the feasibility of generating electrochemical SOFI images of Alexa 647 tagged microtubule filaments in COS-7 cells. A side-by-side comparison of SOFI imaging was performed on the same cell based on UV laser-controlled switching and electrochemically modulated switching of Alexa 647 molecules. The amplitude of the positive electrochemical potential determines the molecule density per frame which was fine-tuned to match the density under UV activation condition for fair comparison. The frequency of oscillating potential was set to frame rate, striking a good balance between the dye refreshing rate and the acquisition speed.

We then applied XC-SOFI, a spatial, temporal, correlation SOFI approach, to run SOFI analysis of SOFI images based on electrochemical or laser control[11]. The XC-SOFI analysis considers pixel correlations in both the spatial and temporal domain that provides several advantages compare to the original temporal correlation approach[10]. It allows for accurate estimation of the image PSF that is useful for subsequent image deconvolution operation. It creates additional virtual pixels between the physical camera pixels so that hardware optical magnification was not required[11]. Upon generating the 2^nd^, 3^rd^ and 4^th^ order SOFI image from 6000 frames for each condition (Figure 2 a, c), significant enhancement was evident in all orders of the SOFI images generated based on electrochemical control compared with the laser-controlled condition. In the SOFI image generated from UV laser activation (Fig. 2a), the tubulin filaments exhibited a markedly discontinuous and punctate appearance. Certain segments of the tubulin fibers were notably absent, as marked by yellow arrow in the zoomed in image, likely attributed to the diminished photoactivation efficiency under UV illumination. The image generated via the electrochemical approach displayed greater smoothness (Fig. 2c), and more details of the sample can be revealed. As indicated by a red arrow in the zoomed in image, the three closely packed tubulin filaments in the bottom left corner in the 4^th^ row could be visually separated in the 4^th^ order SOFI image generated by electrochemical control with good confidence, which was not possible in the UV generated SOFI image. It is worth noting that, under similar image contrast settings, the overall pixel brightness of the 4^th^ order SOFI image generated by electrochemical control was 3 times brighter than its UV counterparts, see Supplementary Figure 1. This can be attributed to the increased correlation under the accelerated switching rate using electrochemical approach.

**Figure 2.**
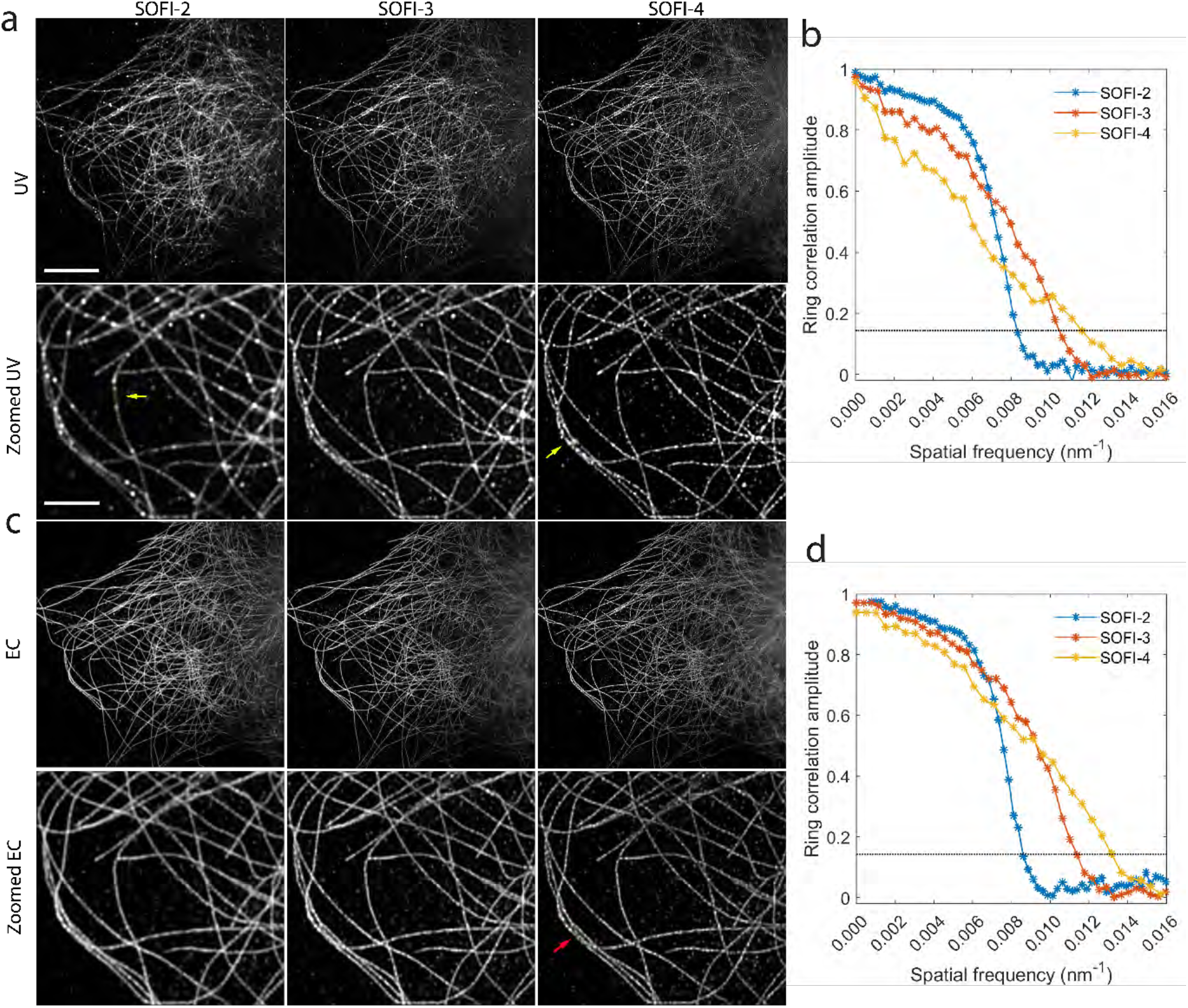
SOFI images of Alexa 647 labelled microtubules under UV (a-b) and electrochemical activation control (c-d). **a**, SOFI-2, SOFI-3 and SOFI-4 is the 2^nd^, 3^rd^ and 4^th^ order SOFI images collected under UV activation condition (0.5 W cm^-2^ of UV laser and 1 kW cm^-2^ 642 nm laser were on). **b**, Fourier ring correlation analysis of the SOFI images shown in a. **c**, The 2^nd^, 3^rd^ and 4^th^ order SOFI images collected under electrochemical activation condition (1 kW cm^-2^ 642 nm laser was on, the electrochemical potential was oscillated between -0.8 V to 0.1 V with a frequency of 20 Hz). **d**, Fourier ring correlation analysis of the SOFI images shown in c. The black dotted line at 0.143 defined the resolution cutoff frequency. The SOFI images displayed were a summation of 6 SOFI images calculated from 6 independent stacks of 1000 frames. Scale bar = 10 µm and 2 µm in the first and second row, respectively.

A rigorous analysis for assessing imaging resolution, Fourier ring correlation (FRC), was used to evaluate the image resolution of the SOFI images[27]. Mathematically, the FRC analysis performs cross-correlation of pixels located inside the increasing concentric rings of the two Fourier transformed images. The intersect of the FRC curve to one seventh of the maximal FRC correlation value at 0.143 was often regarded as a resolution limit of the image[27]. Since we had six 2^nd^ order SOFI images from 6 subset of 1000 frame stacks, we calculated the FRC from two SOFI images generated from even and odd number of the 6 stacks. As the calculated FRC curves of different SOFI order shown in Fig. 2 b and d, as the SOFI order increased, the intersects was right shifted, implying image resolution was increased. More importantly the improvement was more significant with the electrochemical switching method. The resolutions for the 2^nd^, 3^rd^ and 4^th^ SOFI image were 120.5 nm, 86.7 nm and 75.2 nm, respectively for electrochemical based SOFI. In contrast, the resolutions were 122 nm, 96.1 nm and 87.0 nm for 2^nd^, 3^rd^ and 4^th^ order UV based SOFI images. The overall amplitude of the FRC curve was substantially higher at all spatial frequencies in electrochemistry SOFI compared to the UV counterpart. This implies that the image features in the electrochemical controlled SOFI sample were highly consistent across all the length scales. In contrast, the amplitude of the FRC curve from the UV controlled sample decayed rapidly even at low spatial frequencies. The overall lower amplitude of FRC curve in the UV controlled sample implying low consistency of features in the image, which could be caused by large heterogeneity in pixel intensity, and low signal to noise levels in the UV based SOFI image.

### Fast higher order SOFI imaging based on electrochemical switching

One advantage of SOFI imaging, compared to other super-resolution techniques like STORM is that SOFI can handle high molecule density and low signal-to-noise levels in raw images[10, 15, 16]. Therefore, SOFI holds the potential for faster super-resolution imaging compared to localization-based approach[15, 19]. It has been reported recently that low-excitation laser together with extended exposure time are required to optimize the STORM image resolution by reducing emitter density and increase the signal to noise ratio per frame[28], which came at a cost of more than 15 mins acquisition time[29]. Using the electrochemistry modulation, we demonstrated a 3-fold increase of the dye switching rate, which should theoretically enable faster frame rate setting for SOFI imaging[17]. The higher order the SOFI images will improve the resolution, but the differences in dye switching cause excessive pixel brightness scaling in higher order SOFI image that prevent realization of the theoretical SOFI image resolution[10, 15]. In our results in Figure 1 and 2 we showed the 2-fold decrease of heterogeneity of dye switching, which could reduce the pixel scaling effect of the SOFI image. These two key findings motivated us to use electrochemical controlled fast switching of dyes to unlock the full potential of SOFI by taking higher order SOFI images with fast acquisition speed using sCMOS camera[30].

To enhance the switching kinetics of the fluorophores, we applied an oscillating sinusoidal potential between -0.8 V to 0.1 V with a frequency of 20 Hz during SOFI image acquisition. The frame rate for SOFI imaging was set at 100 Hz, and the acquisition time for collecting a data set of 12,000 frames was 124 seconds. For evaluation of the SOFI image quality, we additionally acquired a STORM image of the same cell. To ensure an adequate signal to noise ratio for molecule localization in STORM, the camera exposure time was set to 40 ms per frame. To reduce the emitter overlapping issues within a diffraction area, an electrochemical potential of -0.4 V was applied during STORM data acquisition. It took 13.3 minutes to collect 20,000 frames of images for reconstructing the final STORM image.

We then calculated the 1^st^ to 6^th^ orders of SOFI image, as shown in Figure 3a (SOFI images on whole-cell are provided in Supplementary Figure 2). To reduce the photobleaching effect on SOFI imaging analysis, we performed SOFI analysis from 12 stacks of images (each stack consisting of 1,000 frame) and summed the 12 SOFI images into the final SOFI image. In the 1^st^ order SOFI image, vertical stripes resembling noise patterns were observed due to the high variance in readout noise and different offset values at each column of the sCMOS camera[31] (Fig. 3a first column). However, the noise pattern was completely absent in ≥ 2 order SOFI images. This absence is attributed to the fact that electronic noise does not correlate in time and space. This reconfirmed that ≥ 2 order SOFI image is mostly electronic noise free[11]. It is evident that both the resolution and contrast of the image was significantly improved in ≥ 2nd order SOFI analysis. From the 2^nd^ to the 6^th^ order SOFI image, individual microtubule filaments in the cell become increasingly visible. In the zoomed-in region shown in Figure 3a (second row), two closely spaced tubulin fibers, highlighted by the yellow arrow, could only be resolved in the ≥4th order SOFI image. Importantly, the tubulin filaments are smooth in all order of SOFI image, there was no visible discontinuous and punctate appearance due to pixel scaling in higher order traditional SOFI image[10, 15, 32].

**Figure 3.**
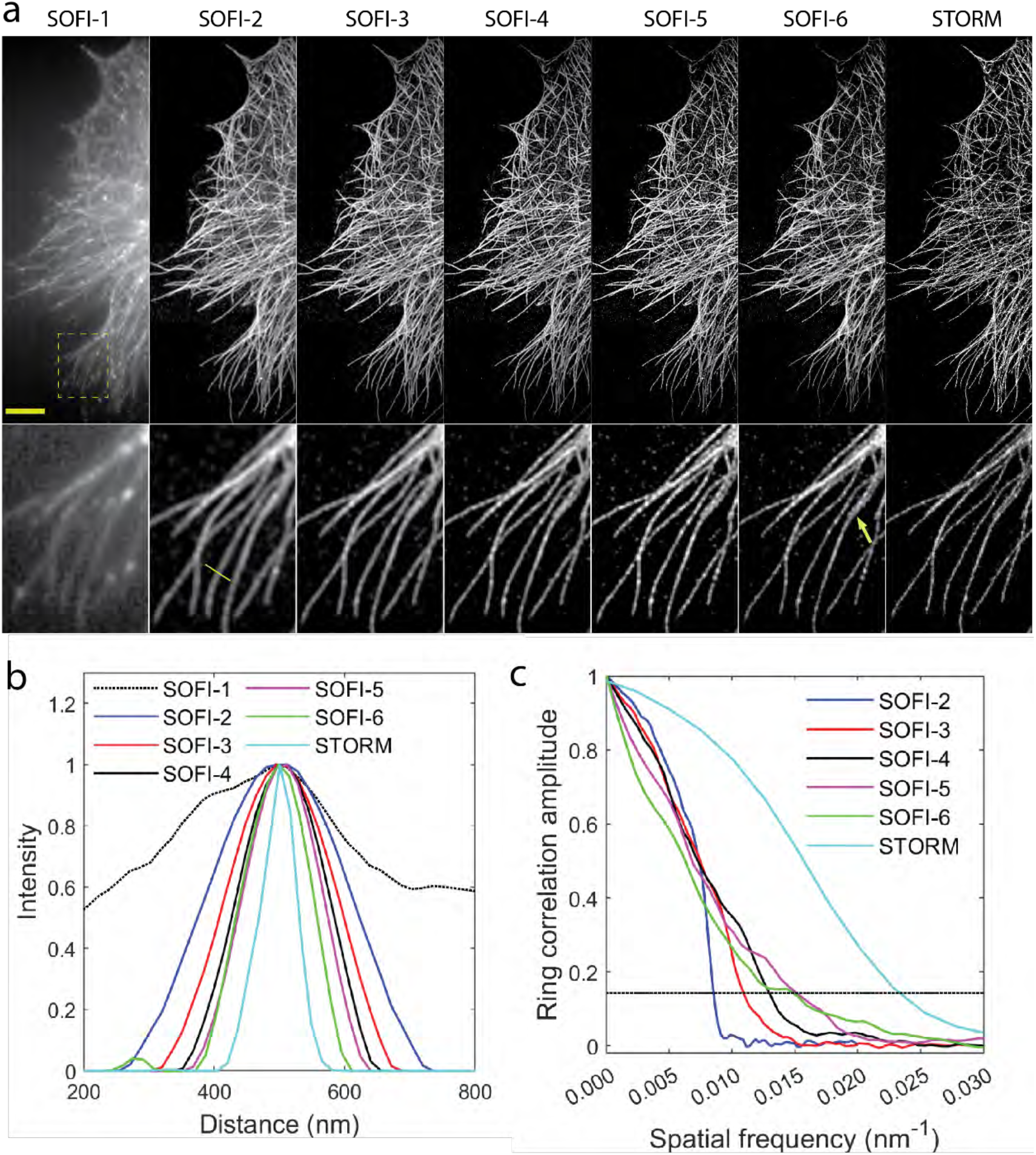
Fast higher order SOFI analysis. **a**, Plot of 1^st^ to 6^th^ order SOFI and STORM images collected from the same cell. A zoomed in region of the yellow dotted box highlighted area was displayed in the second row. Scale bar = 5 µm and 1 µm in the 1^st^ and 2^nd^ row in a, respectively. **b**, FWHM line profile of the yellow line shown for the images displayed in a. **c**, FRC analysis of the images showing in a. The solid lines are smoothed by cubic splines. The abscissa at which the FRC curve intersects with the dotted line represents the resolution limit.

We estimated the image resolution using both full width half maximum (FWHM) and FRC approach. FWHM method extract image resolution directly from line profile of local filament like structure[29, 33], and FRC estimates image resolution systematically of the entire image[33]. By fitting the line profile across a single microtubule fiber (highlighted in yellow line, Figure 3a 2^nd^ row 2^nd^ column) to a Gaussian function, and the FWHM of the fitted Gaussian across the microtubules were 434 ± 47 nm, 140 ± 4 nm, 108 ± 3 nm, 89 ± 2 nm, 78 ± 1 nm, 70 ± 1 nm for the 1^st^ to 6^th^ order SOFI, and 40 ± 1 nm for the STORM image, respectively. FRC analysis showed the whole image resolution for 2^nd^ to the 6^th^ order SOFI images were 117 nm, 91 nm, 78 nm 62 nm and 65 nm, respectively. The STORM image showed the highest image resolution at 43 nm. Although FWHM suggests the microtubule is sharpest in the 6^th^ order SOFI image, the 5^th^ order SOFI image produced the highest resolution according to the FRC analysis. Visually, the higher order SOFI provide comparable details of the sample as in STORM image. In summary, we showed that with fast switching rate of dyes using the electrochemical approach, we had successfully suppressed the discontinuous artifacts in higher order SOFI image, which helped to realize the full potential of higher order SOFI image.

The fast acquisition rate of SOFI demonstrated above has motivated us to explore high throughput super-resolution imaging over large field of view (FOV). One of the ongoing directions in super-resolution imaging in biology is high throughput quantitative imaging that collect large dataset so that the complexity of the biological process can be fully captured where the variability due to noisy biological processes can be averaged out. One way to expand the dimensionality is to perform tile scans over large FOV. Currently, most of the super-resolution imaging techniques are limited to small FOV due to lengthy acquisition time[34]. SOFI stands out as an exception as it requires much smaller number of frames for image reconstruction. Previously, it was demonstrated that not only the switching kinetics but also the density of activated molecules per frame can influence SOFI image resolution[15, 35]. Given that we can easily control molecule density by increase the positive electrochemical potential, we tested SOFI image resolution at different molecule densities as function of number frames used for SOFI calculation. As shown in Fig. 4a-b and Supplementary movie 3 that in general more frames are required to reach the full resolution of higher order SOFI. However, we noticed under higher molecule density, not only less number of frames is needed to reach the best SOFI resolution, but the higher image resolution can also be achieved for the same order SOFI image. For instance, for 2^nd^, 3^rd^ and 4^th^ order SOFI, 400, 700 and 1500 frames are required to reach peak resolution of 150 nm, 100 nm and 90 nm for low density condition, while only 100, 300 and 1100 frame are needed to reach peak resolution of 130 nm, 80 nm and 73 nm for high density condition. At 50 Hz aquation rate, this took 2, 6 and 22 seconds to acquire, respectively. Given the minimal gain in resolution but much lengthier aquation time in 4^th^ order SOFI than the 3^rd^ order SOFI. We prioritized the time resolution to produce only SOFI-2 and SOFI-3 images by taking only 400 frames in each tile position.

**Figure 4.**
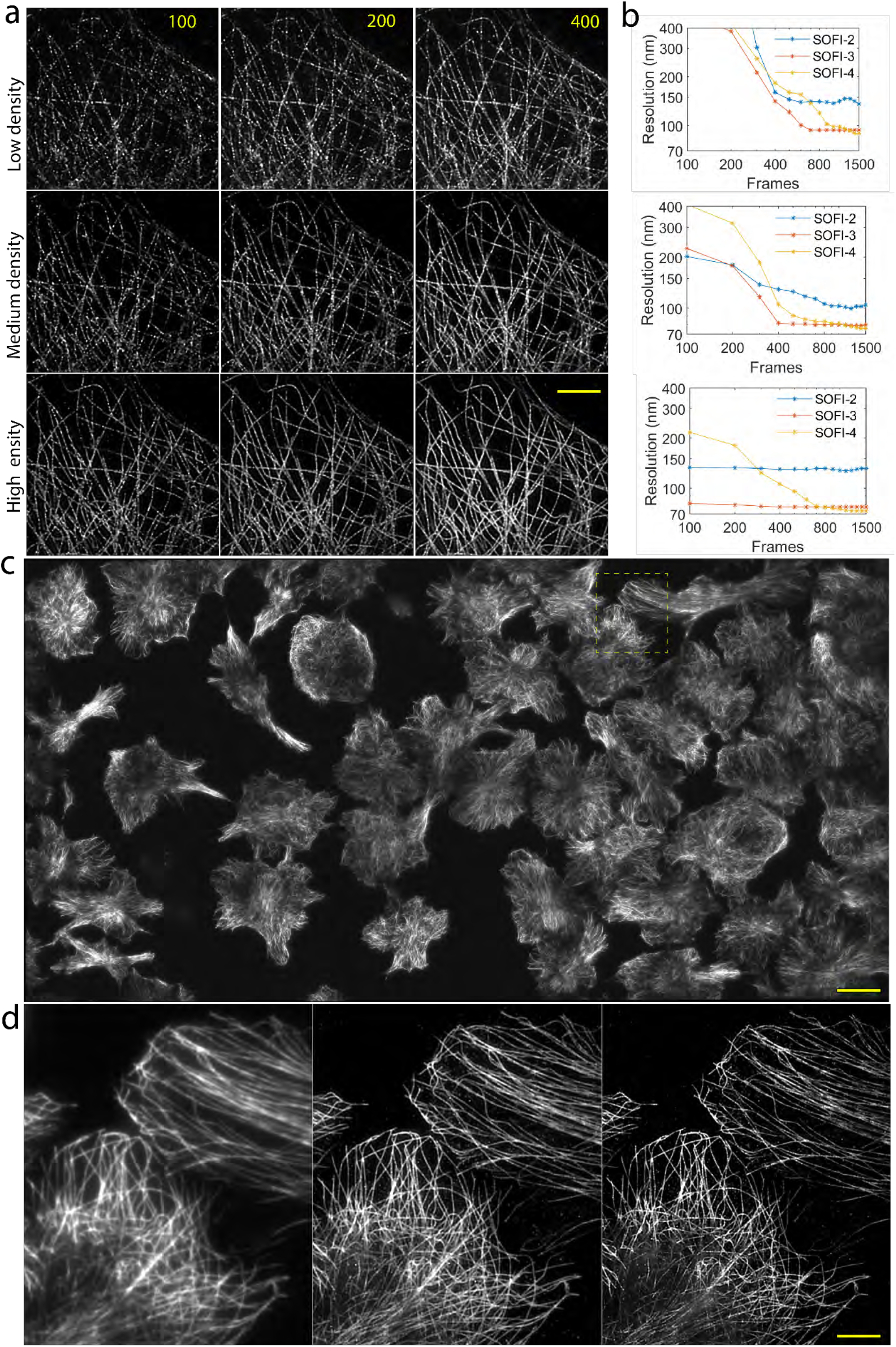
Rapid large FOV super-resolution imaging by SOFI. a, Representative SOFI-3 images generated from indicated number of frames under low (top row), medium (middle row) and high (bottom row) molecule density. b, SOFI-2, SOFI-3 and SOFI-4 image resolution as a function of number of frames used for SOFI calculation. c, Large FOV stitched tile scan TIRF image over area of 0.66 × 0.33 mm. Electrochemically assisted SOFI imaging of the same FOV was acquired subsequently. D. TIRF, SOFI-2 and SOFI-3 images of the zoomed in area of the dotted box highlighted region in c. Full resolution tile scan SOFI2 and SOFI3 image are provided in Supplementary figure 3. Scale bar = 5, 35 and 7 µm in a, c and d, respectively.

We performed an 8 x 4 tiles scanning with a total FOV of 0.66 × 0.33 mm area with a total data acquisition time of 4.5 minutes. Each tile stack was processed separately, and the generated SOFI images of the same order were stitched using a global optimization approach[36], where the maximum cross-correlation of the Fourier transformed image between the pairs produce the best overlaying between the tiles. Over 40 cells were imaged in the FOV as shown in the tile stitched TIRF image shown in Fig. 4c. Distinct improvement of resolution for SOFI-2 and SOFI-3 image can be seen in zoomed in region (Fig. 4d). The full resolution image of whole tile scan SOFI-2 and SOFI-3 image can be seen in Supplementary Figure 3 and Supplementary movie 4. Given the typical acquisition time of 10 minutes for each tile in STORM imaging, electrochemically controlled SOFI image could reduce the acquisition time by 75 folds.

Overall, our work demonstrates enhancing the switching behavior of fluorophore probes with electrochemistry adds significant value for advancing SOFI imaging. Through an external oscillating electrochemical potential with high frequency, we demonstrated the capability for effectively switching the fluorophores on an ITO surface that addressed two key practical limitations of fluorophore switching for high-resolution and fast SOFI imaging. Firstly, we diminished the heterogeneity of pixel brightness in the SOFI image by maintaining uniform dye switching rate across the sample, irrespective of the spatial location and time of data acquisition. This results greatly reduced the sparsity and discontinuity artifacts in a SOFI image. Secondly, we increased the switching rate of Alexa 647 by a factor of 3, enabling increased signal fluctuation in a shorter timeframe. With rapid frame rate using the sCMOS camera, we could produce high quality SOFI image with ∼130 nm resolution in 2 seconds, and 80 nm resolution in 6 seconds. This offers direct advantage of SOFI in large FOV high throughput super-resolution imaging, where we could reduce the acquisition time by a factor of 75 compared to STORM imaging. Electrochemical switching-based SOFI thus presents a robust alternative route for fast high-resolution imaging.

## Methods

### Chemicals and materials

All chemicals, unless noted otherwise, were of analytical grade and used as received from Sigma-Aldrich (Australia). Aqueous solutions were prepared with Milli-Q water of 18.2 MΩ cm resistivity. Through the work, the imaging buffer for SOFI and STORM data collection was oxygen scavenging tris buffer (adjust to pH 8) contains 50 mM Tris, 10 mM NaCl, 10% glucose, 0.5 mg/mL glucose oxidase, 40 μg/mL catalase, and 50 mM cysteamine.

Indium tin oxide (ITO) coated glass coverslips (8-12 Ω, 22×22 cm, SPI Supplies, USA) was utilized to perform electrochemistry and optical imaging at the same time.

Immunofluorescence staining: COS-7 cells (ATCC CRL-1651) was used for immunostaining. The cells were cultured in Dulbecco′s Modified Eagle′s Medium with 10% FBS, penicillin and streptomycin, and incubated at 37°C with 5% CO_2_. COS-7 cells were plated in 6-well plates with ITO slides the day before fixation. The immunostaining procedure for microtubules consisted of: fixation for 7 min at 37°C with 4% paraformaldehyde (Sigma) in PBS, washing with PBS, permeabilization for 5 min with 0.2% Triton X-100 and 3% BSA in PBS, incubation for 1.5 h with rabbit anti α-tubulin monoclonal antibody (ab216650, Abcam) diluted to 2 µg mL^−1^ in blocking buffer (0.2% Triton X-100 and 3% BSA in PBS), washing with PBS, incubation for 30 min with secondary anti-Rabbit IgG antibodies labelled with Alexa 647 at a concentration of ∼2.5 µg mL^−1^ in PBS; washing with PBS, fixation for 5 min with 4% paraformaldehyde in PBS and finally washing with PBS.

### Electrochemistry

Chronoamperometry, and large amplitude sinusoidal voltammetry were performed by SP-200 Potentiostat (Bio-Logic, France). All the electrochemistry was carried out in a custom chamber (Chamlide EC 22, Live Cell Instrument Co., Ltd., Republic of Korea) containing an Ag|AgCl|3M KCl reference electrode and a Pt-mesh counter-electrode, where the working electrode was the ITO coated coverslips.

### Image acquisition

The SOFI and STORM images were collected on Zeiss Elyra 7 Lattice SIM TIRF microscope (Zeiss, Jena, Germany) that coupled with sCMOS camera (edge 4.2, PCO). The collimated and linearly p-polarized 642 nm laser was reflected from the 642 nm long pass dichroic mirror and focused at the back focal plane of the 100 X 1.46 NA Oil objective. The focus is laterally shifted alone the back focal plane to provide either EPI or TIRF illumination. The TIRF angle was identical between the glass and ITO surface. A 1.6 × magnification OptoVar lens was inserted in the detection path to reduce pixel size to 100 nm. The fluorescence was collected by the same objective and guided to a cooled sCMOS camera.

For electrochemical SOFI imaging, 1 kW cm^-2^ 642 nm laser was used for excitation. For UV assisted SOFI imaging, 1 kW cm^-2^ 642 nm laser was used for excitation and 5-50 W cm^-2^ 405 nm laser was on for photoactivation. The camera exposure time were set to 10-30 ms as indicated in the text.

For STORM imaging, 2 kW cm^-2^ 642 nm laser was used for excitation and 5-50 W cm^-2^ 405 nm laser was applied for photoactivation. The camera exposure time was 30 ms. In tile scan large FOV SOFI imaging, the images were acquired in TIRF mode under oscillating electric potential of 20 Hz between -0.8 V to 0.1 V. The camera was set in crop mode with 950 x 950 format as the illumination intensity drops off near the edge area at full pixel resolution of 1280 X 1280. 32 stacks of 1000 frames were collected for each tile position with 12% overlay between the tiles. Each tile was processed separately to generate individual SOFI images. The ImageJ stitching plugin[36] was used for stitching of the tiles to provide the best overlaying between the tiles. Essentially, the maximum cross-correlation between the Fourier transformed neighbouring images gives the lateral shift needed to generate best overlaying between the image pairs.

### Temporal pixel correlation analysis

The image stacks collected under UV activation condition (UV, 0.5 kW cm^-2^ of 642 nm laser and 5 W cm^-2^ of UV laser are on) or electrochemically controlled condition (EC, the potentials were oscillated between -0.8 V and 0.4 V with a frequency of 10 Hz, 5 kW cm^-2^ of 642 nm laser is on) were applied for pixel autocorrelation analysis as an estimation of the average on time of the dyes. The intensity over time of individual pixel was defined as the raw correlation time trace. The correlation was calculated as *G*(τ) = ⟨*F*(*t*)*F*(*t* + τ)⟩/⟨*F*(*t*)⟩⟨*F*(*t* + τ)⟩.

where *F* is the pixel intensity, *t* is pixel acquisition time, ***τ***is correlation time, <> stand for averaging. The correlation curves from all the pixel contained molecules were averaged to produce the averaged correlation. The variance across those the pixels were used to analyze the heterogeneity of dye switching in Figure 1d.

### SOFI image analysis

XC-SOFI image analysis approach was used to generate SOFI images[11, 15, 37]. A previous published custom written Matlab XC-SOFI code was used to generate the SOFI image (GitHub - lob-epfl/sofitool). Briefly, the spatial-temporal cross cumulant between adjacent pixels were calculated to generate a grid of virtual pixels between true physical pixels. For higher order of cumulant analysis, a finer grid of virtual pixels at the geometrical centre of a set of selected pixels were generated. There were different combination sets of pixels could generate virtual pixels at the same spatial locations. In this case, those cumulant values from different set were averaged. Depend on the distance of the virtual pixels from the true physical pixels, a Gaussian shaped distance weighting factor was applied to correct the cumulant amplitude of the virtual pixels. This is because the correlation amplitude decreases gradually at increasing spatial lags. From the generated raw SOFI image, the PSF of the resulting image was estimated from the profile of the spatial correlation analysis[38], which was used in the subsequent deconvolution process[11]. As a result, the final resolution of the n^th^ order SOFI image was increased by a factor of n instead of 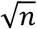 as in the original SOFI image[10]. The raw image data was split into 1000 frame stacks, and SOFI calculation were performed on each stack. The final SOFI image was summation of all generated SOFI images.

### STORM image analysis

The STORM image analysis was performed on the Zeiss Zen PALM software. Briefly, the local intensity maxima of each frame were identified by thresholding with a signal to noise ratio of 6. Once the coarse position of the molecules was identified. A circular region with a diameter of 7 pixels was segmented as isolated diffraction spot to fit to a 2D Gaussian function. The peak position of the Gaussian curve as the representation of the spatial coordinate of the molecule would be estimated at high accuracy upon fitting. Reasonably separated fluorophores with minimal overlapping were accounted by interactive fitting, subtracting and refitting procedure[39]. No grouping was applied to regroup the switching events from the same molecule as no molecular quantification was needed in current work. A model based automatic drift correction provided in the Zen software was used for post processed image drift correction. To generate STORM image, the Zen generated STORM table was loaded into a custom written Matlab algorithm. A binary image with pixel size 20 times finer than the original image was generated with the spatial coordinates of all fluorophores assigned to 1. A 2D Gaussian function with sigma of 20 nm was convoluted with the binary image to generate the rendered STORM image.

### Image resolution analysis

The FWHM analysis was used as an alternative simple approach for direct image resolution estimation. The line profile *f*(*x*) over a structure smaller than the diffraction limit was fitted to the one-dimensional Gaussian function 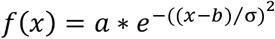. Where, *x* is pixel position, *a, b* and σ is the amplitude, centre and width of Gaussian function, which is used as the FWHM value in current study. The FRC analysis were utilized as resolution evaluation method[27]. Briefly, two images of the same sample were Fourier transformed using the *fft2* inbuild function in Matlab. The real part of the Fourier transformed image was shifted with zero frequency at the image centre by the *fftshift* function. For n^th^ ordered SOFI image, the pixel size was reduced by a factor of √*n*, which set the maximal spatial frequency in the Fourier transformed image. To perform cross correlation of pixels at various spatial frequency, the images were cropped by a ring-shaped mask with width of 7 pixels that centred at the zero frequency. The diameter of the ring was increased from 14 pixels until the inner edge of the ring reach the edge of the Fourier transformed image. The cross correlation of pixels within the two rings was normalized by the standard deviation of pixels within each ring.

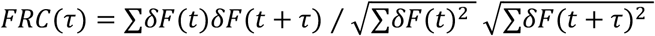

Where *δF* is the deviation of the pixel intensity from the mean intensity within the ring, ∑stands for sum, ***τ***is the spatial lag from the zero frequency in the image center. To reduce the edge effect in the Fourier transformed image, a vertical and horizontal line strip with width of 4 pixels were used to crop out those pixels prior to the FRC correlation analysis. For FRC analysis of the STORM image, the molecule coordinates were randomly split into two groups, and two separate STORM images were generated from the divided molecule coordinates. For the FRC analysis of SOFI image, FRC from a pairs of SOFI images produced from the even and odd image stacks were calculated.

## Acknowledgement

The authors gratefully acknowledge J. F. Berengut for provision of the DNA origami sample. The authors also acknowledge technical assistant by the Katharina Gaus Light Microscopy Facility, University of New South Wales. J.J.G. acknowledges the Australian Research Council Discovery Grant Program (DP220103024) and a National Health and Medical Research Council Investigator Grant (GNT1196648). Y.M. acknowledges the Earlier Career Fellowship funding from the National Health and Medical Research Council of Australia (APP1139003).

## Notes

### Competing Interest Statement

The authors have declared no competing interest.

