## Supplementary Information for "Electrochemically controlled switching of dyes for enhanced super-resolution optical fluctuation imaging (SOFI)"

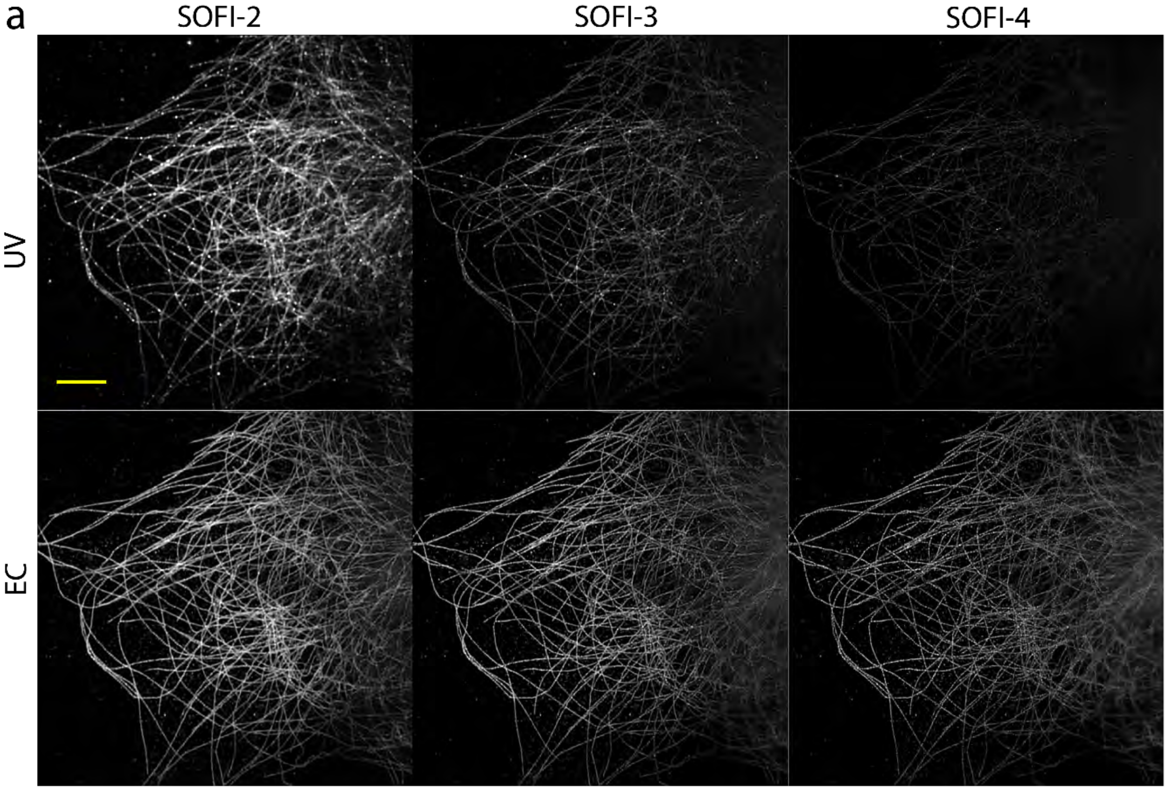


**Supplementary Figure 1.** a, 2[nd], 3[rd] and 4[th] order SOFI images produced by UV illumination (1[st] row, 0.5 W cm$^{-2}$ of UV laser and 1 kW cm$^{-2}$ 642 nm laser were on) or by electrochemical modulation (2[nd] row, 1 kW cm$^{-2}$ 642 nm laser was on, the electrochemical potential was oscillating between -0.8 V to 0.1 V with a frequency of 20 Hz) under the same image contrast settings. The SOFI images displayed were a summation of 6 SOFI images calculated from 6 independent stacks of 1000 frames. Scale bar = 2 μm.

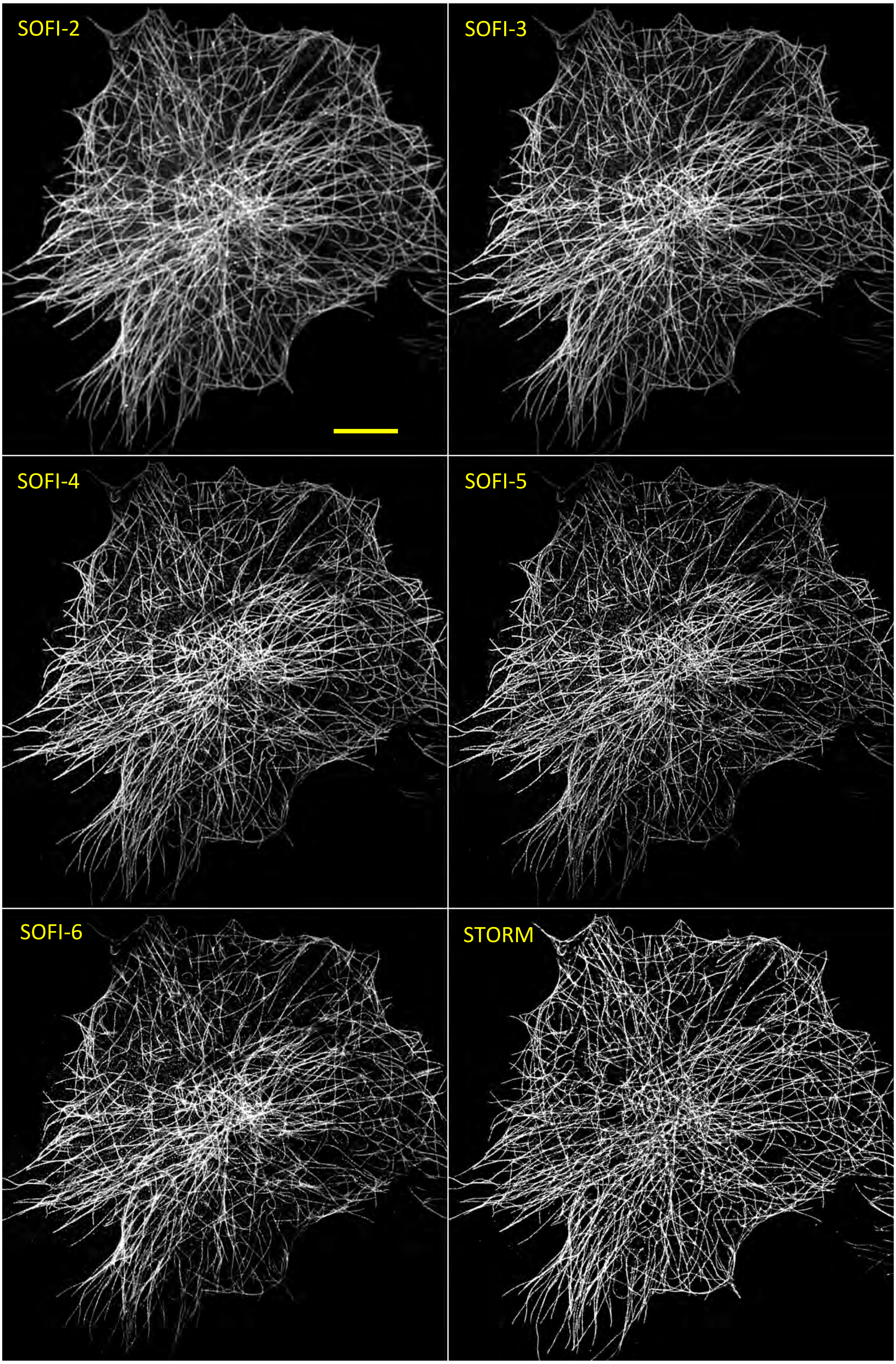
SOFI-2
SOFI-3
SOFI-4
SOFI-5
SOFI-6
STORM

**Supplementary Figure 2.** Comparison for 2nd to 6th order SOFI and STORM image of microtubule in the same COS-7 cell. For SOFI imaging, 1 kW $cm^{-2}$ 642 nm laser was on, the electrochemical potential was oscillated between -0.8 V to 0.1 V with a frequency of 20 Hz. For STORM, , 2 kW $cm^{-2}$ 642 nm laser was used for excitation and 5-50 W $cm^{-2}$ 405 nm laser was applied for photoactivation. Scale bar = 5 µm.

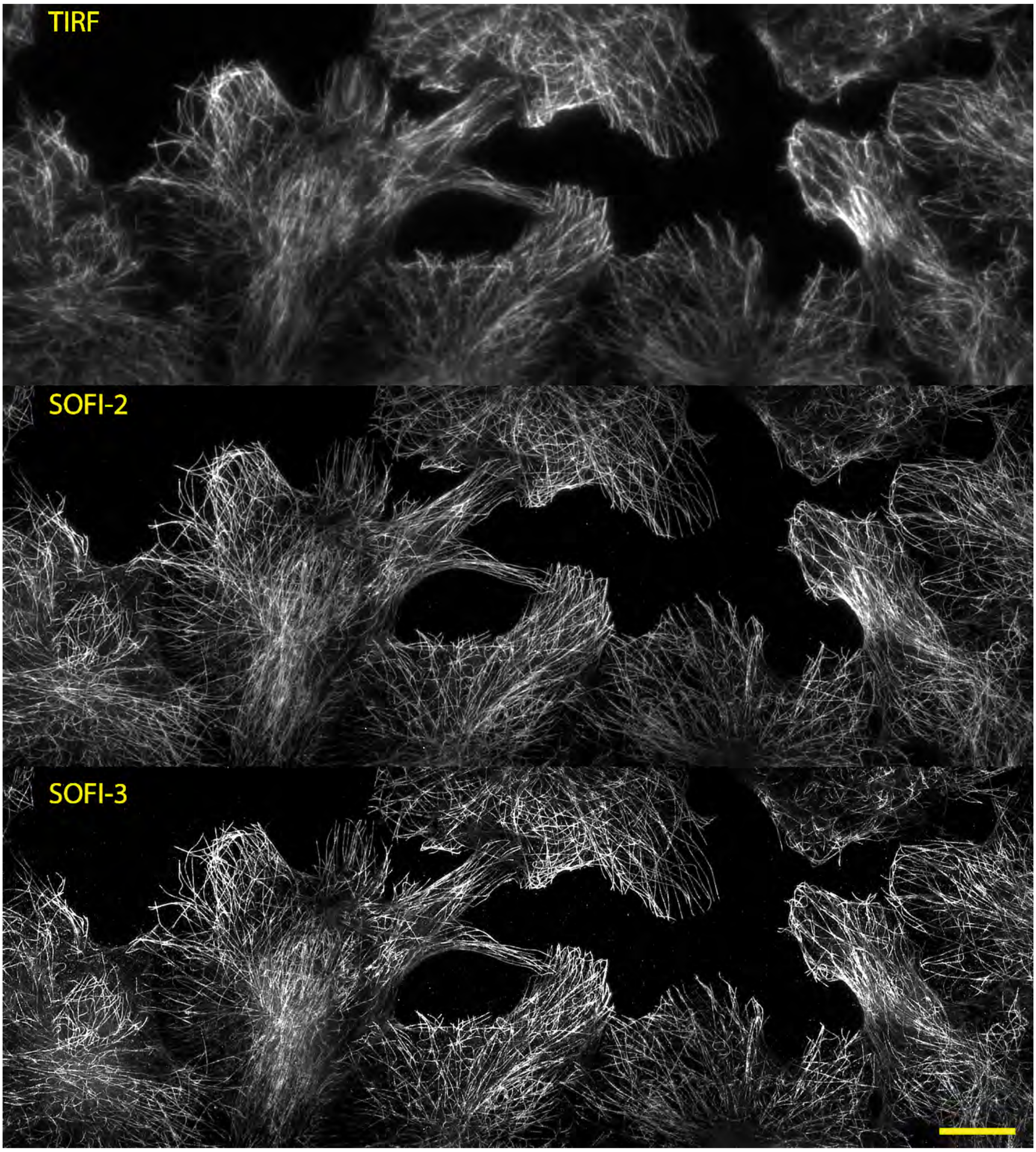


**Supplementary Figure 3.** A representative cropped area of large FOV tile scan of TIRF (top), 2nd and 3rd order SOFI images of microtubules in COS-7 cells generated under electrochemical modulation condition (1 kW $cm^{-2}$ 642 nm laser was on, the electrochemical potential was oscillated between -0.8 V to 0.1 V with a frequency of 20 Hz). Scale bar = 10 µm.

**Supplementary movie 1**. Reversible electrochemical switching of single Alexa 647 labelled origami on ITO coverslip in dSTORM buffer. The electrochemical potential was alternating at 1 Hz frequency between -0.8 V and 0.4 V. Images were acquired in TIRF mode. The camera exposure time was 50 ms.

**Supplementary movie** 2. SOFI images of the same sample with Alexa 647 labelled microtubles under either UV activation of and electrochemcial modulated switching conditions. Left, under UV activation condition, 0.5 W $cm^{-2}$ of UV laser and 1 kW $cm^{-2}$ 642 nm laser were on. Right, under electrochemical modulation condition, 1 kW $cm^{-2}$ 642 nm laser was on, the electrochemical potential was oscillated between -0.8 V to 0.1 V with a frequency of 20 Hz. The moelcular density per frame was comparable under UV or electrochemical conditions.

**Supplementary movie 3**. The image resolution as a function of frames used for SOFI calculation in low, medium and high molecular density conditions. The frame number used are indicated. From left to right column, the images were $2^{nd}$ , $3^{rd}$ and $4^{th}$ order SOFI images, respectively.

**Supplementary movie 4**. Moving window display of TIRF and $2^{nd}$ order SOFI images of the large FOV tile scan of Alexa 647 labelled microtuble in COS-7 cells.
